# Microbiota stability in healthy individuals after single-dose lactulose challenge – a randomized controlled study

**DOI:** 10.1101/424531

**Authors:** Sandra Y. Wotzka, Markus Kreuzer, Lisa Maier, Mirjam Zünd, Markus Schlumberger, Bidong Nguyen, Mark Fox, Daniel Pohl, Henriette Heinrich, Gerhard Rogler, Luc Biedermann, Michael Scharl, Shinichi Sunagawa, Wolf-Dietrich Hardt, Benjamin Misselwitz

**Affiliations:** Institute of Microbiology, D-BIOL, ETH Zürich, CH-8093 Zürich, Switzerland; Abdominal Center: Gastroenterology, St. Claraspital, Kleinriehenstrasse 30, 4068 Basel, Switzerland; Department of Gastroenterology, University Hospital Zurich (USZ), and Zurich University, Rämistrasse 100, 8091 Zurich, Switzerland

**Author notes:** Contributed equally to this paper (Sandra Y. Wotzka, Markus Kreuzer, Lisa Maier, Mirjam Zünd, Shinichi Sunagawa, Wolf-Dietrich Hardt, Benjamin Misselwitz). **Clinical Trials. Gov registration number:** NCT02397512. **Correspondence:** Prof. Dr. Wolf-Dietrich Hardt Institute of Microbiology, D-BIOL ETH Zürich Vladimir-Prelog-Weg 4 CH-8093 Zürich, Switzerland, Prof. Dr. Shinichi Sunagawa Institute of Microbiology, D-BIOL ETH Zürich Vladimir-Prelog-Weg 4 CH-8093 Zürich, Switzerland, Dr. med. Benjamin Misselwitz Department of Gastroenterology University Hospital Zurich (USZ), and Zurich University Rämistrasse 100 CH-8091 Zurich, Switzerland.

**Keywords:** Microbiota, Lactulose, Hydrogen, Salmonella typhimurium, Escherichia coli, Constipation, Hepatic Encephalopathy

## Abstract

**Background and aims:** Lactulose is a common food ingredient and widely used as a treatment for constipation or hepatic encephalopathy and a substrate for hydrogen breath tests. Lactulose is fermented by the colon microbiota resulting in the production of hydrogen (H_2_). H_2_ is a substrate for enteropathogens including *Salmonella* Typhimurium (*S*. Typhimurium) and increased H_2_ production upon lactulose ingestion might favor the growth of H_2_-consuming enteropathogens. We aimed to analyze effects of single-dose lactulose ingestion on the growth of intrinsic *Escherichia coli* (*E. coli*), which can be efficiently quantified by plating and which share most metabolic requirements with *S*. Typhimurium.

**Methods:** 32 healthy volunteers (18 females, 14 males) were recruited. Participants were randomized for single-dose ingestion of 50 g lactulose or 50 g sucrose (controls). After ingestion, H_2_ in expiratory air and symptoms were recorded. Stool samples were acquired at days −1, 1 and 14. We analyzed 16S microbiota composition and abundance and characteristics of *E*. *coli* isolates.

**Results:** Lactulose ingestion resulted in diarrhea in 14/17 individuals. In 14/17 individuals, H_2_-levels in expiratory air increased by ≥20 ppm within 3 hours after lactulose challenge. H_2_-levels correlated with the number of defecations within 6 hours. *E. coli* was detectable in feces of all subjects (2 x 10^2^ - 10^9^ CFU/g). However, the number of *E*. *coli* colony forming units (CFU) on selective media did not differ between any time point before or after challenge with sucrose or lactulose. The microbiota composition also remained stable upon lactulose exposure.

**Conclusion:** Ingestion of a single dose of 50 g lactulose does not significantly alter *E*. *coli* density in stool samples of healthy volunteers. 50 g lactulose therefore seems unlikely to sufficiently alter growth conditions in the intestine for a significant predisposition to infection with H_2_-consuming enteropathogens such as *S*. Typhimurium (www.clinicaltrials.gov NCT02397512).

## Introduction

Humans coexist with trillions of microbes on their body surfaces, collectively referred to as microbiota. One of the key benefits these microbes confer to their host is colonization resistance (CR), i.e. protection against invasion and infection by pathogens. It is conceivable that during gut colonization, the pathogen has to compete with the resident intestinal microbiota for nutrients and binding sites (nutrient-niche hypothesis) (1, 2). Yet, the mechanistic details underlying the protective effect by the microbiota have not been fully elucidated.

Several perturbations can disrupt colonization resistance. Antibiotic treatment can disturb the intestinal microbiota, predisposing the host to infections with enteric pathogens including *Salmonella* Typhimurium (*S*. Typhimurium) and *Clostridium difficile* in humans (3) and in a murine model (1, 4, 5). Likewise, according to the nutrient-niche hypothesis, providing the ecosystem with an additional nutrient should also open new niches, which might reduce colonization resistance for certain pathogens. Effects of long-term and short-term (< 1 day) perturbations might thereby differ since long-lasting interventions will trigger complex secondary responses of the gut ecosystem.

The disaccharide lactulose is used as a food ingredient (named galacto-fructose) (6, 7) and as a prescription or over-the-counter medication for constipation (8) or hepatic encephalopathy in individuals with liver cirrhosis (9). The lactulose breath test uses lactulose to diagnose small intestinal bacterial overgrowth (10). As lactulose cannot be absorbed during the passage through the host’s small intestine (11), it reaches the large intestine where it enriches the nutrient pool available to the gut microbiota. After arrival, lactulose is fermented by certain gut microbiota members resulting in the production of short chain fatty acids (SCFA), hydrogen (H_2_) and methane (12, 13). The production of these compounds explains the laxative properties and also the side effects (abdominal pain and bloating) of this drug. Based on these observations, it seems likely that the intake of lactulose might significantly alter nutrient availability in the colon and thereby change the colonic microbiota composition. Effects of short-term lactulose exposure on the gut microbiota have been insufficiently studied.

We hypothesized that lactulose may also promote gut luminal growth of *Enterobacteriaceae* like *S*. Typhimurium or *Escherichia coli* (*E. coli*). *E. coli* shares most metabolic requirements of S. Typhimurium and growth of *E. coli* in stool samples is considered a surrogate for favorable growth conditions of enteropathogens such as *S*. Typhimurium (14). In contrast to *S*. Typhimurium, *E. coli* can grow on lactulose, i.e. by taking up and degrading this disaccharide. However, *E. coli* might also benefit directly or indirectly from the sugar monomers galactose and fructose, which are released upon degradation of lactulose by certain members of the gut microbiota (15, 16) or by fermentation products like H_2_. H_2_ can serve as an electron donor for *S*. Typhimurium and other *Enterobacteriaceae* (17–21). Data in our murine model of *S*. Typhimurium colitis demonstrated significant attenuation of a *S*. Typhimurium mutant deficient in H_2_-metabolism during initial stages of infection (18). H_2_ therefore constitutes a critical metabolic intermediate exploited by *S*. Typhimurium for efficient colonization of the large intestine (18).

Short-term exposure to lactulose might therefore be a model for a disease mechanism with a defined manipulation of the intestinal environment, potentially altering the microbiota composition and affecting susceptibility to enteric infections. Thus, we hypothesized that the intake of lactulose might further promote gut luminal growth of *Enterobacteriaceae* such as *E. coli* and *S*. Typhimurium. These effects might manifest rapidly since previous studies demonstrated alterations in microbiota composition within 24 hours after dietary interventions (22–25). Thereby, a single lactulose dose will allow monitoring direct effects of this intervention, avoiding the complexity of secondary physiological and microbiological compensatory responses upon long-term exposure.

In this study, we aimed at monitoring short-term effects of lactulose intake on human volunteers upon a single lactulose dose. We tested for preferential growth of *Enterobacteriae* (enterobacterial blooms) which would indicate increased risk for infection by *Enterobacteriaceae* in healthy individuals and by extrapolation in patients.

## Experimental Procedures

### Sample size calculation

To test the primary hypothesis of our study, that lactulose exposure would increase growth of
*E. coli* (as a marker for *Enterobacteriae*) the following sample size calculation was performed: To be clinically relevant, the increase in *E. coli* counts between day −1 and day 1 should be at least one order of magnitude. In a conservative calculation, we estimated the logs of the differences in *E. coli* counts between day −1 and 1 to be normally distributed with a wide standard deviation of two orders of magnitude. In a one-tailed paired analysis, a sample size of 27 and 36 individuals would be required to detect an increase of *E. coli* counts by one order of magnitude with a power of 80% or 90%, respectively.

### Study participants

For this study, 32 healthy volunteers were recruited via advertising. Exclusion criteria were a history of intestinal surgery (except hernia repair, appendectomy or anorectal surgery), pregnancy, relevant gastrointestinal symptoms (i.e., symptoms that cannot be ignored), medical or psychiatric conditions requiring ongoing management or intake of medication affecting gastrointestinal function (for instance laxatives, opioids, non-steroidal anti-inflammatory drugs, proton pump inhibitors or antibiotics within the last 4 weeks). To ensure proper handling of stool samples only employees of ETH Zurich or Zurich University, familiar with freezing and storing samples at −80°C were recruited. All participants provided informed consent and were compensated for their expenses. The first participant was included on the 30.01.2015, the last participant completed the study on 30.04.2015. No follow up was performed.

**Figure 1:**
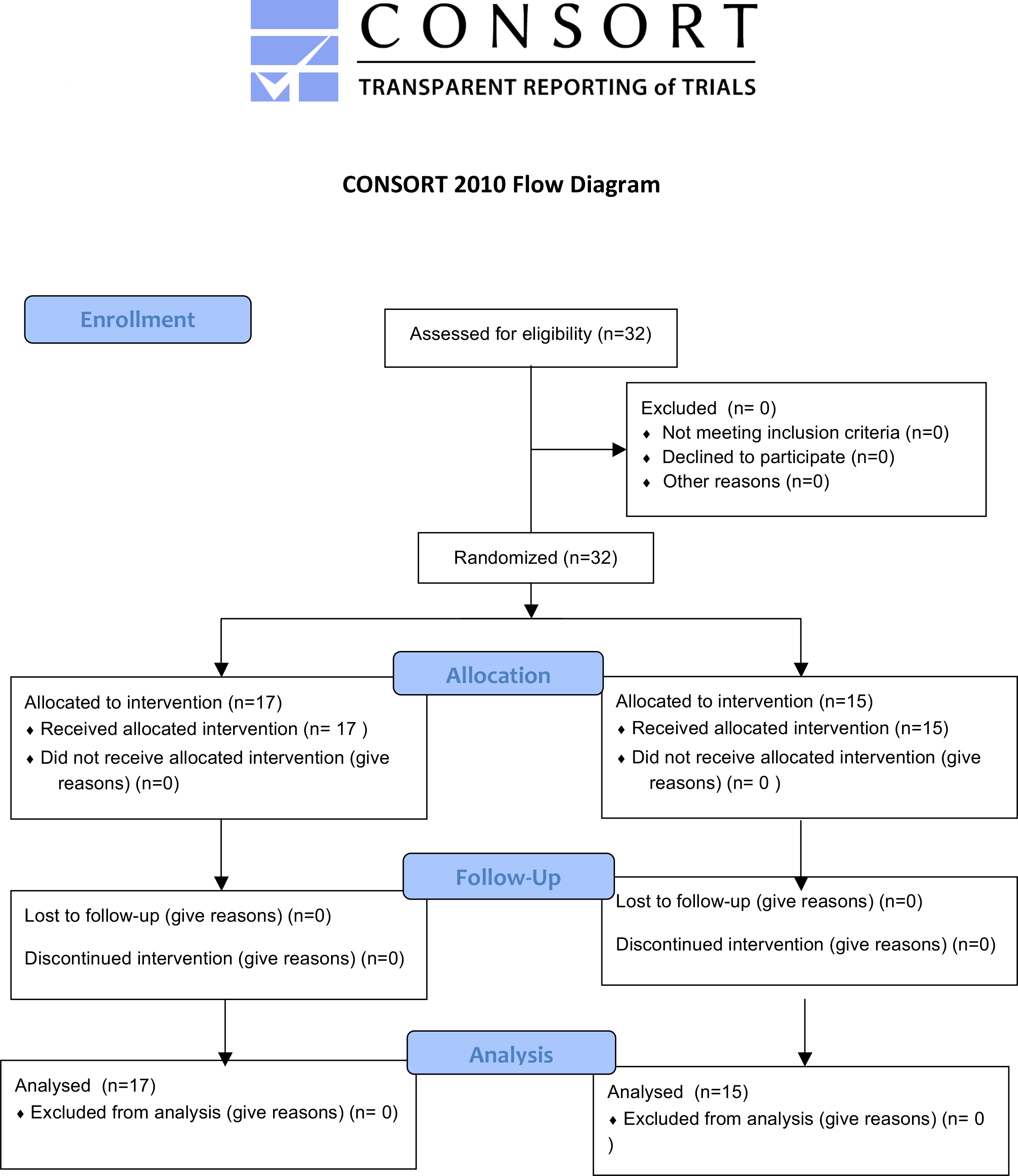
Consort Flow chart

Participants were screened during an interview and epidemiological data, information regarding gastrointestinal function or symptoms, medical history, medication, gastrointestinal function and life style (vegan, vegetarian or Western diet, sport [hours of exercise per week], alcohol consumption, smoking) were recorded (Table 1). Before the intervention day, participants fasted for 12h with only water allowed prior to the intervention.

**Table 1:** Epidemiological and clinical characteristics of the study population. BSS: Bristol stool scale; IQR: interquartile range.

|  | **Sucrose** | **Lactulose** |
| --- | --- | --- |
| **Number of participants** | 15 | 17 |
| **Age**<br>median (IQR; range) | 25 (23-28.5; 21-36) | 24 (22-31; 21-47) |
| **Gender** female n/N (%)<br>male n/N (%) | 9/15 (60 %)<br>6/15 (40 %) | 9/17 (53 %)<br>8/17 (47 %) |
| **Stool frequency**<br>every 2-3 days n/N (%)<br>1x per day n/N (%)<br>2-3x per day n/N (%) | 2/15 (13 %)<br>10/15 (67 %)<br>3/15 (20 %) | 3/17 (18 %)<br>12/17 (70 %)<br>2/17 (12 %) |
| **Stool consistency (BSS)**<br>median (IQR; range) | 4 (3-4; 3-7) | 4 (3-4; 3-4) |
| **Diet**<br>Western diet<br>Vegetarian diet | 15/15 (100 %)<br>- | 15/17 (88 %)<br>2/17 (12 %) |
| **Non-smokers** n/N (%) | 15/15 (100 %) | 17/17 (100 %)<sup>a</sup> |
| **Physical exercise per week (hours)**<br>median (IQR; range) | 4 (3-5.5; 2-8) | 3 (2-5; 0-12) |
| **Medication**<br>Yes n/N (%)<br>No n/N (%) | 8/15 (53 %)<sup>b</sup><br>7/15 (47 %) | 7/17 (41 %)<sup>c</sup><br>10/17 (59 %) |
| **Ethnicity**<br>Caucasian n/N (%)<br>Non-Caucasian n/N (%) | 14/15 (93 %)<br>1/15 (7 %)<sup>d</sup> | 14/17 (82 %)<br>3/17 (18 %)<sup>e,f,g</sup> |

<sup>a</sup> Minimal nicotine consumption in one person (3-4 cigarettes/week)
<sup>b</sup> 3x hormonal contraception, 1x thyroid hormone replacement, 3x ibuprofen
<sup>c</sup> 3x hormonal contraception, 1each ibuprofen, paracetamol, thyroid hormone replacement
<sup>d</sup> born in Vietnam, living in Switzerland for 2 years
<sup>e</sup> parents from Sri Lanka, participant born in Switzerland
<sup>f</sup> born in Turkey, living in Switzerland for 4 years
<sup>g</sup> born in Cameroon, living in Switzerland for 3 years

Our study was approved by the Ethics committee of Zurich county (KEK-ZH 2014-0358) and registered at clinicaltrials.gov (NCT02397512).

### Study design

The participants were randomly assigned to two groups (http://www.graphpad.com/quickcalcs/randomize1.cfm) using the sealed envelope method and treated in a double blinded fashion. The control group (n=15) received a sham treatment of 50 g sucrose (Migros, Zurich) dissolved in 200 ml water. The intervention group (n=17) consumed a drink containing 50 g lactulose (Legendal 12g, Zambon, the Netherlands) dissolved in 200 ml water (Figure 2). H_2_ levels in parts per million (ppm) in expiratory air of participants were measured at 0 min (baseline), 15 min, 30 min, 60 min, 90 min, 120 min, and 180 min after the consumption of the respective sugar solution using the HydroCheck breath measuring device (Neomed Medizintechnik GmbH Üchtelhausen, Germany) (Figure 2). Methane content of expiratory air was recorded using Breath Collection Kit (BreathTracker SC, QuinTron USA). Due to technical issues, methane measurements could be performed only in the last 14 individuals included in our study. At all time points participants were asked to evaluate the strength of nausea, bloating, pain, stool alterations and borborygmi (bowel sounds) using a 4-point scale (1 - no symptoms, 2 - light symptoms which can be ignored, 3-marked symptoms which cannot be ignored but would not affect daily activity and 4 - severe symptoms affecting daily activities. An additional questionnaire recorded bowel consistency (maximum Bristol stool scale, BSS) and frequency of bowel movements during 6 hours after the consumption of the respective sugar solution.

**Figure 2:**
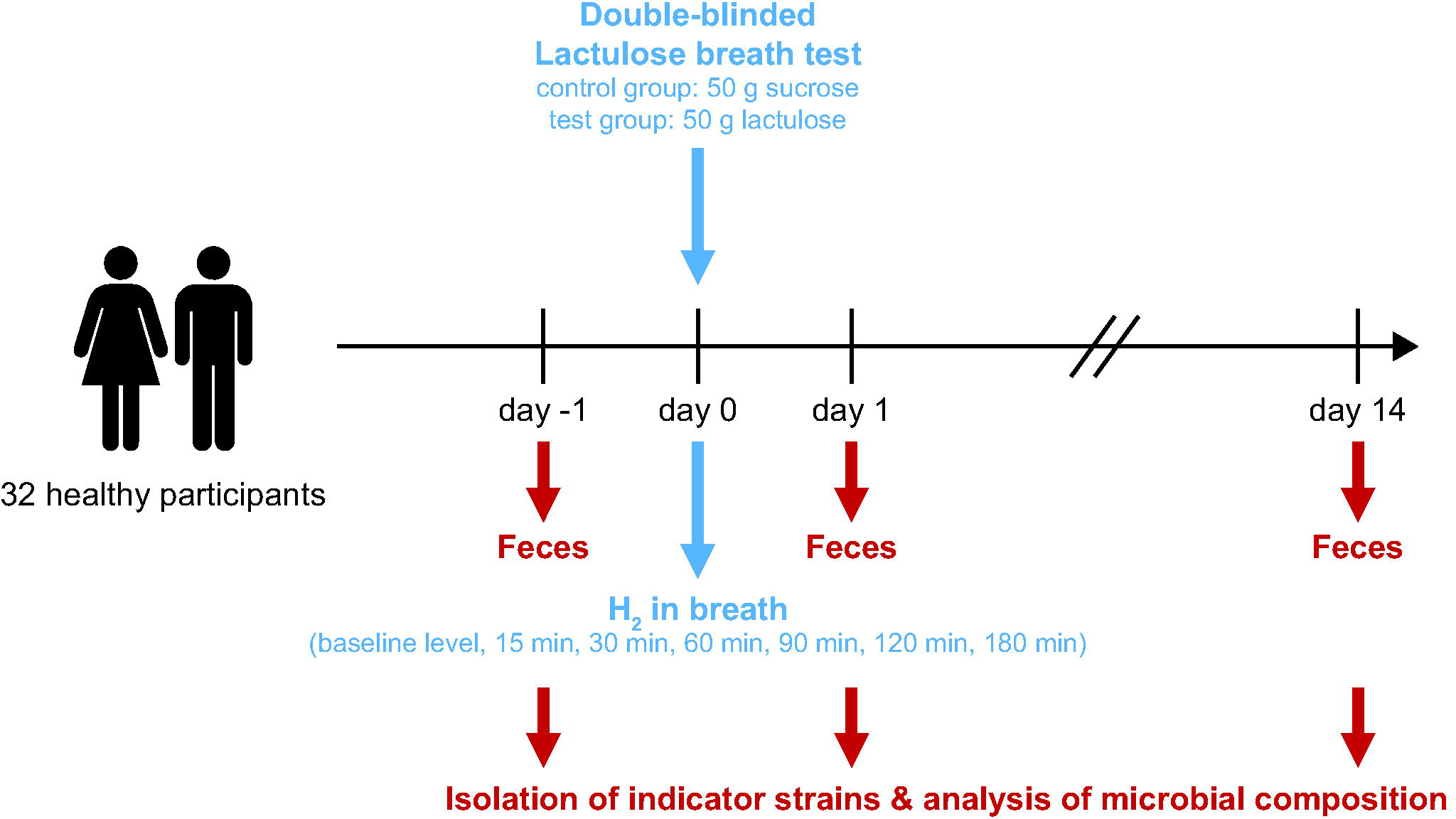
Overview of the design of our study. 32 healthy participants were randomly assigned to either the control group (sucrose) or to the experimental group (lactulose). Fecal samples were collected one day before treatment, one day after treatment and two weeks after treatment for further microbial analysis. On the day of treatment, hydrogen levels were measured in expiratory air at the indicated time points.

To assess effects on the resident gut microbiota, fecal samples were collected one day before, one day after, and two weeks after lactulose or sucrose exposure (Figure 2). Half of the fecal sample was stored in a test tube containing trypticase soy broth (Oxoid/ Thermo Fisher Diagnostics AG, Pratteln, Switzerland) with 15% (w/v) glycerol (Sigma-Aldrich, Buchs, Switzerland) (26). This sample was used for the analysis of the colony forming units (CFU) per gram feces and the isolation of single bacterial strains. The other half of the fecal sample was stored in a test tube without any supplements and used for the analysis of microbiota composition. Within one hour after collection, samples were delivered by the participants and stored in a −80°C freezer provided by the investigators. To ensure proper handling of samples only individuals familiar with handling of −80°C freezers had been recruited (see inclusion criteria above) and importance of rapid sample transfer was emphasized to all individuals. Timely transfer of the fecal sample to the freezer within 30-60 min was confirmed by questionnaires to participants.

### Microbiota quantification and isolation of *Enterobacteriacae* by agar plating

Fecal samples were transferred to Eppendorf tubes containing 500 µl of phosphate buffered saline (PBS) with tergitol (Sigma-Aldrich) and a sterile metal ball, weighed, and homogenized (1 min at 25 Hz, using a TissueLyser (Qiagen, Hombrechtikon, Switzerland). Differential plating on MacConkey agar plates (Oxoid) without any antibiotics allowed identification of bacterial population size (CFU; i.e. *E. coli*, *Salmonella* spp. and other *Enterobacteriaceae*) within the individual samples. In an attempt to capture the enterobacteriaceal diversity, several morphologically different colonies from each time point were picked and restreaked three times on new MacConkey agar plates. After restreaking, pure single colonies were picked and cultured in lysogeny broth (LB) for 10-12h at 37°C shaking (160 rpm). Bacterial cultures were stocked in peptone glycerol broth (peptone: Oxoid, glycerol: Sigma-Aldrich), shock frozen and stored at −80°C.

### 16S rRNA gene analysis of isolated *Enterobacteriacae* strains

For each bacterial strain, 50 µl of an overnight culture were centrifuged (5min, 8000 rpm), suspended in 100 µl sterile H_2_O and incubated at 99°C for 3 min. Genomic DNA was collected from supernatant after centrifugation (3min, 8000 rpm). 8 µl DNA was amplified by PCR in a 100 µl reaction mixture containing 10 µl buffer (5 PRIME PCR master kit), 5 U Taq polymerase (5 PRIME PCR master kit), 8 µl 2.5 mM dNTPs, 53 µl ddH_2_O and 10 µl of each forward primer (fD1: 5’-AGAGTTTGATCCTGGCTCAG-3’) (27) and reverse primer (rP1: 5’-ACGGTTACCTTGTTAGCACTT-3’) (28) diluted 1:10 in ddH_2_O. The amplification program consisted of an initial denaturation step of 5 min at 94°C; followed by 35 cycles of 1 min at 94°C, 1 min at 43°C, and 2 min at 72°C; and a final extension of 7 min at 72°C. Correct PCR product size was confirmed using agarose gel electrophoresis and ethidium bromide staining, DNA fragments were excised under UV light and extracted using the Wizard SV Gel and PCR Clean-Up System (Promega, Dübendorf, Switzerland).

Excised DNA fragments were sent for sequencing at Microsynth AG Balgach, Switzerland. The sequences were annotated by the basic local alignment search tool (BLAST) (29) for taxonomical identification. Entries were sorted by “Max score” and the top record used to annotate the genus and, if reliable, also the species classification for each bacterial strain. The sequence alignment tool MUSCLE (30) and SplitsTree 4.13 (31) were used to construct phylogenetic trees based on the BLAST annotations.

### DNA extraction for 16S gene-based microbiota composition analysis

DNA was extracted using the QIAamp Stool Mini Kit (Qiagen) with the following changes in the disruption and homogenization steps: 700µl of Buffer ASL, 2 spoons of small glass beads (0.1mm; Bio Spec products) and 1 spoon of large glass beads (0.5-0.75mm; Schieritz&Hauenstein, Laufen, Switzerland) were added to each stool sample. The samples were mixed using the tissue lyser (Qiagen) (3 min, 30Hz) and heated for 5 min at 95°C. Next, samples were again mixed using the tissue lyser (3 min, 30Hz), centrifuged at full speed for 1 min and the supernatants were transferred to a new 2 ml microcentrifuge tube. The pellet was suspended in 200 µl of Lysis Buffer (20mg/ml lysozyme (Sigma-Aldrich); 20mM Tris⋅HCl, pH 8.0; 2mM EDTA; 1.2% Triton) and incubated at 37°C for 30 min. Afterwards, 500 µl of Buffer ASL were added to each sample followed by mixing using the tissue lyser (3min, 30Hz) and heating the suspension for 5 min at 95°C. All samples were mixed again using the tissue lyser (3vmin, 30Hz) and centrifuged at full speed for 1 min. The supernatants were transferred to the same 2 ml tube than used before. Half of an InhibitEX tablet (provided in Kit) was added to each sample and tubes were vortexed immediately until the tablet was completely suspended. Next, samples were incubated for 1 min at room temperature, centrifuged at full speed for 3 min and supernatants were transferred into a new 1.5 ml microcentrifuge tube. The sample were again centrifuged at full speed for 3 min. Meanwhile, 35 μl of proteinase K (provided in Kit) was added to a new 2 ml tube and supernatants from previous centrifugation step were transferred to the 2 ml tube containing the proteinase K. 200 μl of Buffer AL were added and samples were vortexed for 15 sec and incubated at 70°C for 10 min. The next steps were performed according to the manual instructions. The final elution step was performed using 100 μl Buffer AE (pre-heated to 70°C) added to the membrane, then samples were incubated for 1 min at room temperature and centrifuged at full speed for 1 min to elute DNA.

### Library preparation and sequencing for microbiota composition analysis

The 16S rRNA gene libraries were produced using the NEXTflex^®^ 16S V4 Amplicon-Seq Kit 2.0 (Barcodes 1- 96; Bioo Scientific, Austin, Texas, USA). The input concentration of genomic DNA was adjusted to 30 ng/µl for each PCR reaction. The library preparation was performed following the manufacturer’s instructions with the differences that the reaction volume was reduced to 25 µl and a modified primer pair was used for the first PCR reaction. Instead of the original primer pair 515f-806r, the degenerative primer 515F (5′-GTG**Y**CAGCMGCCGCGGTAA-3′) and 806rB (5’-GGACTAC**N**VGGGTWTCTAAT-3’), as described in (32, 33) were used. The first PCR reaction was performed using Q5 High-Fidelity DNA Polymerase (BioConcept, NEB, Allschwil, Switzerland) under the following cycling conditions: 1) initial denaturation: 95°C for 4 min; 2) denaturation: 95°C for 30 sec; 3) annealing: 56°C for 30 sec; 4) extension: 72°C for 90 sec; 5) final extension: 72°C for 4 min. Cycles 2-4 were repeated 8 times. After each reaction, the PCR products were cleaned up using AMPure XP magnetic beads (Beckman Coulter SA, Nyon, Switzerland). The clean PCR products were eluted in 20 µl of resuspension buffer and used in a second PCR reaction for producing multiplexed samples, respectively. The quantity of the amplicons was measured by a Qubit 3.0 fluorometer (Thermo Fisher Scientific). The quality of the amplicons was evaluated using a Fragment Analyzer (Advanced Analytical, Ankeny, IA, USA). The length of the barcoded amplicons was approximately 450 bp. The amplicons were pooled at equimolar concentrations and diluted to a final concentration of 60 ng DNA per 20 µl before loading on the Illumina MiSeq platform. Sequencing was performed at 2 x 250 bp read length at the Functional Genomics Center Zurich.

### Microbiota composition analysis

Initial analyses were performed using USEARCH (version 9.1.13) using custom scripts that performed the following steps: paired reads were merged and quality-filtered using the *fastq_mergepairs* command with default settings. Merged reads were filtered using the *fastq_filter* command (-fastq_maxee 1.0) and only merged reads with perfect primer matches and a minimum length of 100 bp were selected. Sequences were de-replicated using the *fastx_uniques* and clustered into operational taxonomic units (OTUs) at 97% with chimera removal using the *cluster_otus* command (-minsize 2). OTU abundances for each sample were quantified using the *usearch_global* command (-strand both; -id 0.97). Taxonomic annotation was performed by matching OTU sequences against the SILVA database (version 128) using the *usearch_global* command (-id 0.90; -maxaccepts 20; -maxrejects 500; -strand both; -top_hits_only).

Further analyses were performed in RStudio (version 1.0.143 based on [R] version 3.3.3) using the libraries vegan and ggplot2. Read counts were rarefied to 19 000 reads per sample and Shannon’s diversity index calculated using the *diversity* function. For performing a Principal Coordinate Analysis and for calculating intra-individual dissimilarity (grouped by time point or individual), Bray-Curtis distances were computed between pairs of samples using the *vegdist* function. Significant differences between time points for each treatment group as well as phlym-level taxonomic abundances were calculated by paired Wilcoxon tests with FDR correction (0.05) for multiple testing.

Sources of microbial composition variance was tested by permutational (N=999) MANOVA using the *adonis* function.

The exact Mann-Whitney U test, the Spearman R correlation and the Two-way ANOVA were performed using GraphPad Prism Version 7.02 La Jolla, CA, USA. The sample size calculation was performed using G*Power Version 3.1.9.2 (34). A p-value of less than 0.05 was considered significant.

### Antibiotic susceptibility testing

Test antibiotics were chosen according to known genetic determinants of resistance (35). Test tubes containing 2 ml LB media were inoculated with the isolated bacterial strains and incubated for 5h at 37°C shaking (160 rpm). Optical density (OD) 600 was measured and a fraction of the culture was used to inoculate new test tubes containing 2 ml LB media to obtain a final OD 600 of approximately 0.02. Liquid cultures were diluted 1:10 in sterile ddH_2_O and streaked with a cotton swab three times with 45° shifts on Mueller-Hinton agar to obtain a bacterial lawn of a density according to British Society for Antimicrobial Chemotherapy (BSAC) standards (36). Antibiotic discs were placed on the agar using sterile tweezers, followed by incubation at 37°C for 18-20h. The diameter of the zone of inhibition was measured for each antibiotic and bacterial strain were as susceptible, intermediate, and resistant according to the Clinical and Laboratory Standards Institute (CLSI) (37).

## Results

### Study population

Our study recruited 32 healthy volunteers (18 out of 32, 56% female) of young age (median 25, IQR: 22.75-30, Table 1). All individuals had normal bowel habits (frequency of bowel movements every 2-3 days to 2-3 times per day, median BSS 4), were non-smokers and most individuals followed a Western diet with regular physical activity. Medical history did not reveal relevant medical conditions or gastrointestinal complaints; medication included only analgetics, thyroid hormone replacements and hormonal contraception (Table 1). Thus, we recruited a homogenous healthy study population.

### Symptoms upon sucrose and lactulose ingestion

Individuals were randomized with 17 individuals in the intervention group with lactulose ingestion and 15 individuals in the control group with sucrose ingestion, respectively (Figure 2). Following ingestion of the respective disaccharide, 14/17 (82%) of individuals in the lactulose group but only 3/15 (20%) of individuals in the control group reported physical symptoms. Within three hour of observation on a five-point scale, scores for bloating, diarrhea, borborygmi (bowel sounds) and abdominal pain of the lactulose group were significantly higher compared to the sucrose group (bloating: median 2 [range 1-4] vs. 1 [1-2], p=0.0006; diarrhea: median 1 [range 1-5] vs. 1 [1-1], p=0.0128; borborygmi: median 2 [range 1-4] vs. 1 [1-2], p=0.0005; abdominal pain: median 1 [range 1-3] vs. 1 [1-2], p=0.0163; Figure 3, Table S1).

**Figure 3:**
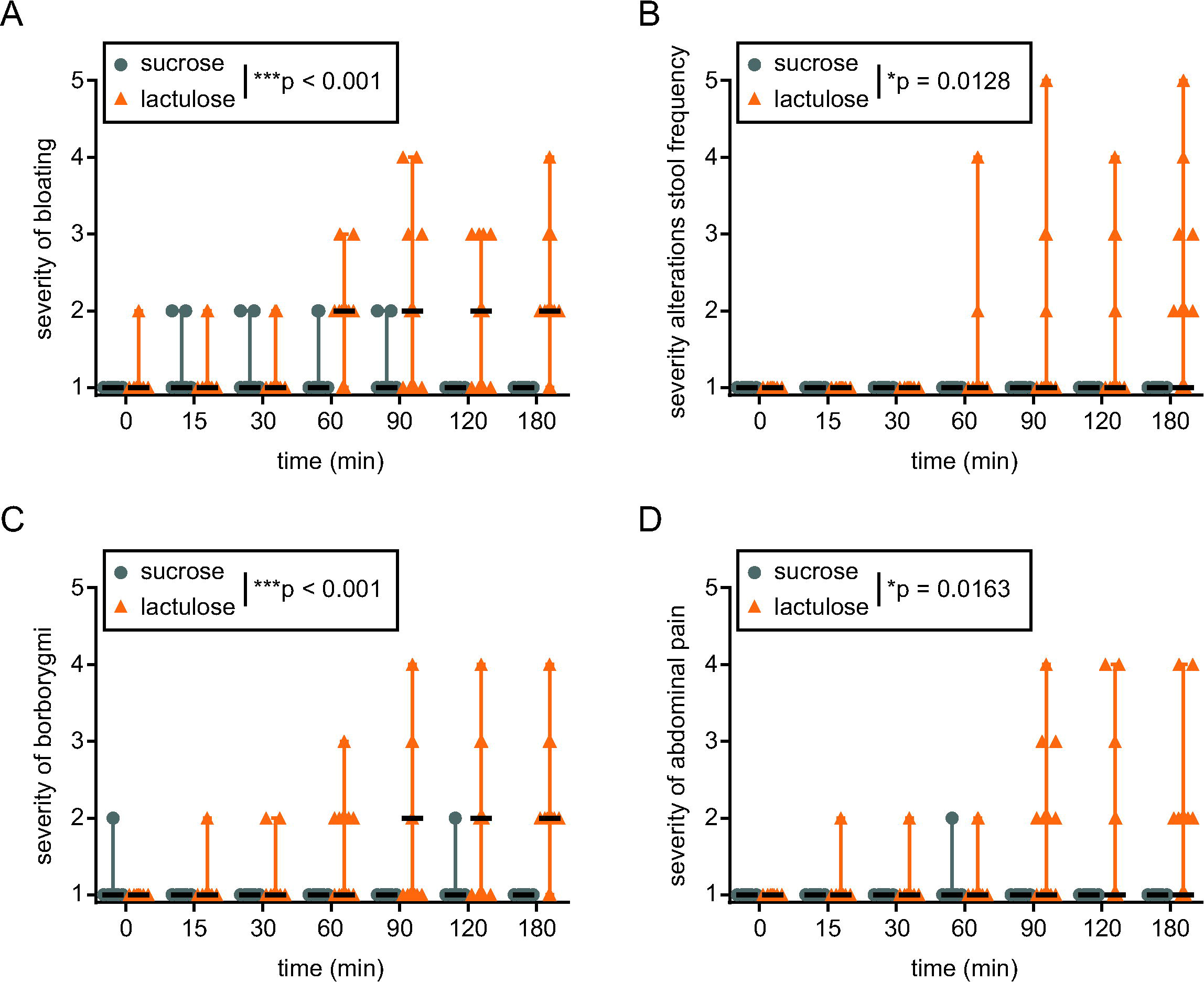
Symptoms upon lactulose or sucrose ingestion. Severity of (A) bloating, (B) stool alterations, (C) borborygmi and (D) abdominal pain was rated by individual participants according to the following scale: 1 - no symptoms, 2 – symptoms which can be ignored), 3 – symptoms which cannot be ignored, but don’t affect daily activities, 4 – symptoms affecting daily activities, 5 – symptoms dominating and severely inhibiting daily activities. *p<0.05, ***p<0.001; Two-way ANOVA with repeated measurements.

On separate questioning, 13 out of 17 individuals reported diarrhea (defined as ≥3 bowel movements within 6 hours after lactulose intake); diarrhea was accompanied by loose stool consistency (Bristol stool scale, BSS ≥5 in all individuals with diarrhea). 5 of those reported ≥4 bowel movements with BSS 7. The remaining 4 out of 17 individuals did not respond to lactulose treatment with no defecation during 6 hours in 3 participants and 1 defecation BSS 5 in the remaining participant. No diarrhea was observed in the control (sucrose) group (data not shown).

After lactulose exposure, H_2_ levels in expiratory air gradually increased for the next 90 min and remained at this level for most individuals until the end of the observation period (180 min, Figure 4). H_2_ levels varied widely (0-146 ppm); for three individuals of the lactulose intervention group, H_2_ levels in expiratory air never increased for more than 20 ppm over baseline; these individuals are by definition H_2_ non-producers (10).

**Figure 4.**
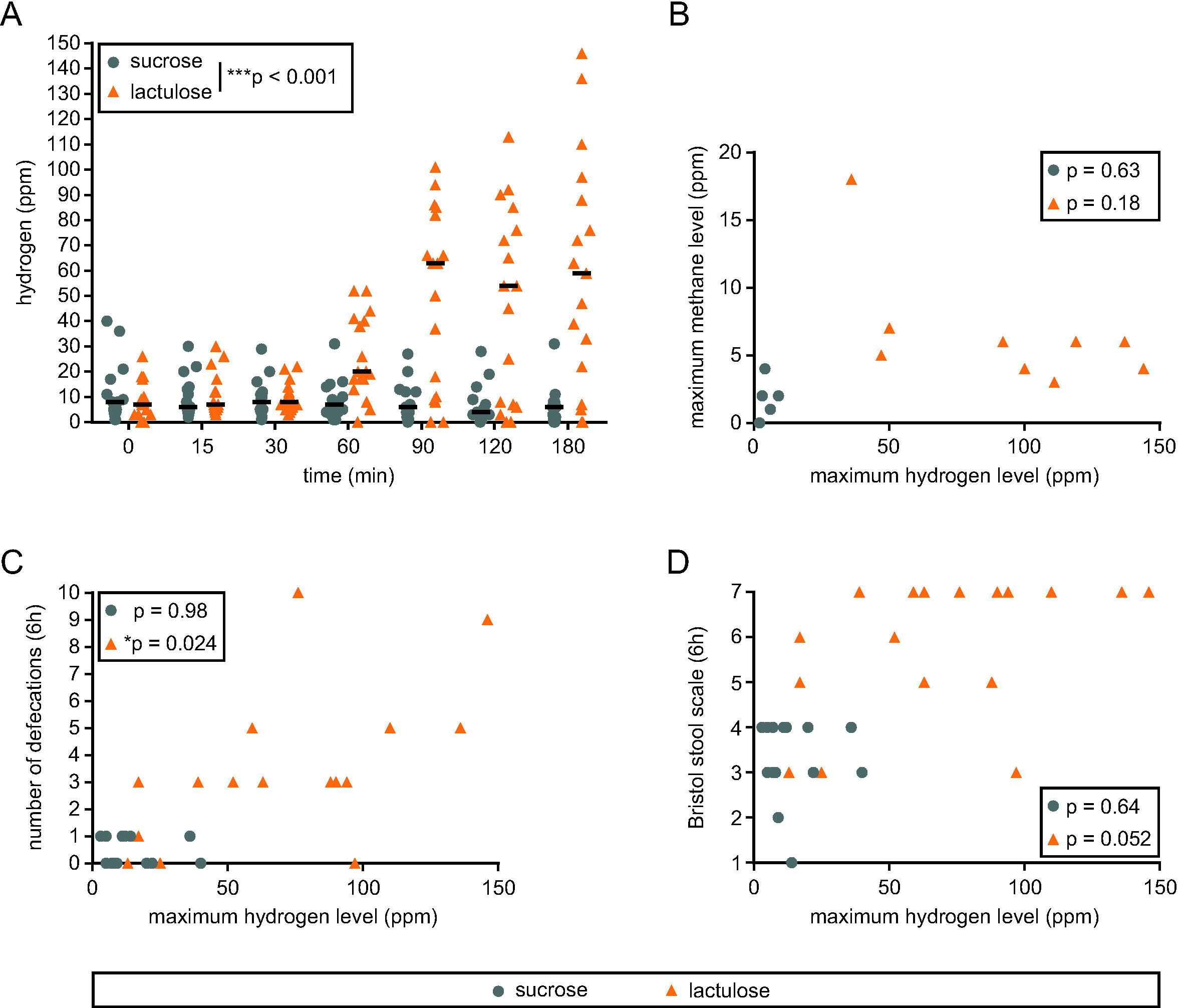
Hydrogen and methane levels and symptoms following lactulose ingestion. (A) Hydrogen levels in parts per million (ppm) following lactulose or sucrose (control) ingestion. ***p < 0.001; two-way ANOVA. (B) Correlation between maximum (max) hydrogen and methane levels. (C) Correlation between maximum hydrogen levels and number of defecations during 6 hours following lactulose or sucrose ingestion. (D) Correlation between maximum hydrogen levels (max) and stool consistency (Bristol stool scale) following lactulose or sucrose ingestion. Spearman R correlation.

For 14 out of 32 individuals (5 in the sucrose group, 9 in the lactulose group) methane (CH_4_) measurements were also taken (Figure 4B); for technical reasons, only the last 14 individuals included into our study could be tested. However, only a single individual had increased CH_4_ levels at baseline (15 ppm) and for no individual CH_4_ levels exceeded 10 ppm over baseline. For the three H_2_ non-producers no CH_4_ measurements were available.

H_2_ levels in expiratory air correlated with number of defecations (p=0.024, Figure 4C); correlation analysis with stool consistency (BSS) revealed a non-significant trend (p=0.052, Figure 4D), suggesting that the intestinal response to lactulose was associated with H_2_ production.

### Bacterial growth in stool samples of lactulose and sucrose treated subjects

We tested whether lactulose ingestion and exposure of the intestine to excess H_2_ increases levels of *E*. *coli* and related *Enterobacteriaceae* in stool samples. Consistency of collected stool samples (as judged by in inspection of collected material) was similar at all time points (day −1, day 1 and day 14), arguing against significant confounding by sample dilution. Stool samples were plated on selective media to quantify *E*. *coli* one day before (baseline), one day after, and two weeks after disaccharide exposure (Figure 5). At all time points, no difference in bacterial numbers could be detected (Figure 5A-C). Variation within the two groups was high, with bacterial densities ranging from 3 x 10^2^ to 10^9^ colony forming units (CFU) per g fecal sample in the control group and 2 x 10^2^ to 8 x 10^8^ CFU per g in the lactulose group (Figure 5A-C).

**Figure 5:**
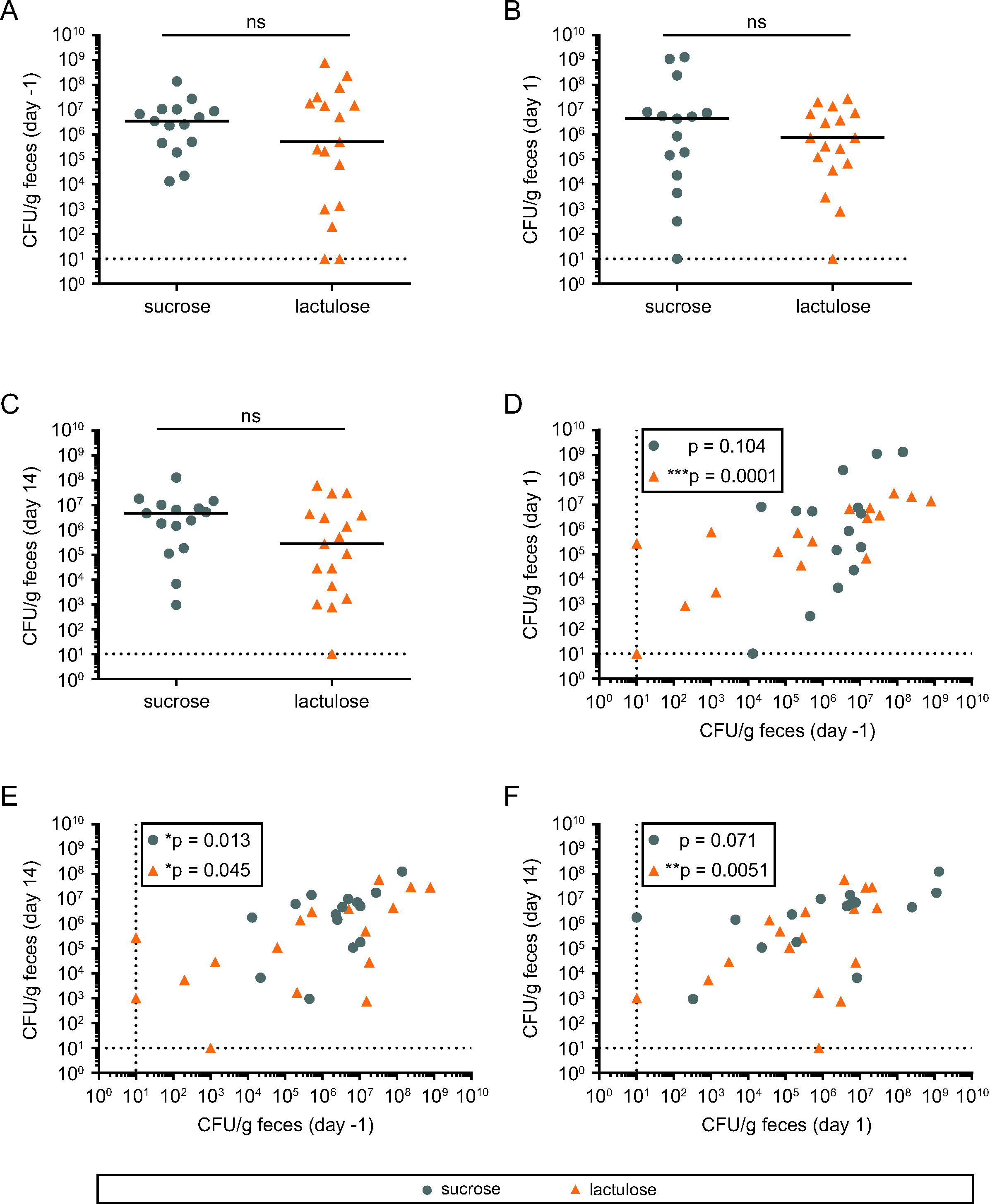
*E. coli* counts in stool samples at different time points. (A-C) Stool samples were plated on selective agar without antibiotics to quantify *E. coli* counts (CFU). ns = not significant; Mann-Whitney U test. (D-F) Correlations of *E. coli* counts within the feces collected at different time points. ns: not significant; *p < 0.05 = significant; **p < 0.01; ****p < 0.001; Spearman R correlation. Dashed lines = detection limit.

CFU counts remained stable over time and no difference for time points could be detected in a paired analysis (p>0.05 for all comparisons, Wilcoxon matched pairs signed-rank test). We found highly significant correlations of the number of CFUs at baseline, day 1 and day 14 in the group with lactulose treatment (Figure 5D-F), indicating stability of *E*. *coli* population size over time and upon lactulose perturbation.

As expected, *E. coli* counts were high, illustrating the abundance of *E*. *coli* in the microbiota of healthy individuals. Considering the similar H_2_ metabolism in *E. coli* and related organisms (e.g. *S*. Typhimurium, *Yersiniae*, *Campylobacter*, *Shigellae* (17–21) our data argue against a pronounced effect of increased H_2_ levels on the growth of potential invading enteropathogens. In line with this interpretation, the number of CFU at the day after lactulose exposure (or the ratio of CFU between time points) did not correlate with maximum H_2_ levels in expiratory air at the intervention day (Figure S1A).

Number of CFUs or ratios of CFUs between days also did not correlate with number of defecations after lactulose exposure (Figure S1B). We noted a positive correlation between CFU and stool consistency (BSS) and the ratio of CFUs on day −1: day 1 in the control group but not in the lactulose group (Figure S1C).

No *Salmonella enterica* colonies were detected in any of the stool samples, as judged from the absence of color-less colonies on the MacConkey agar plates.

### Phylogenetic analysis of stool isolates

To confirm species affiliation of isolated *Enterobacteriaceae*, bacterial strains from 30 participants at one day after treatment were analyzed by sequencing the 16S rRNA gene. Sequences were classified using the basic local alignment search tool (BLAST). All bacteria isolated from MacConkey agar were found to belong to the family of *Enterobacteriaceae* with a large cluster of 20 *E*. *coli* related species (20 *E*. *coli*, 5 *E*. *coli*/ *Shigella*) and a distantly related cluster of 2 *Klebsiella*, 2 *Enterobacter* and 1 *Citrobacter* species (Figure 6).

**Figure 6:**
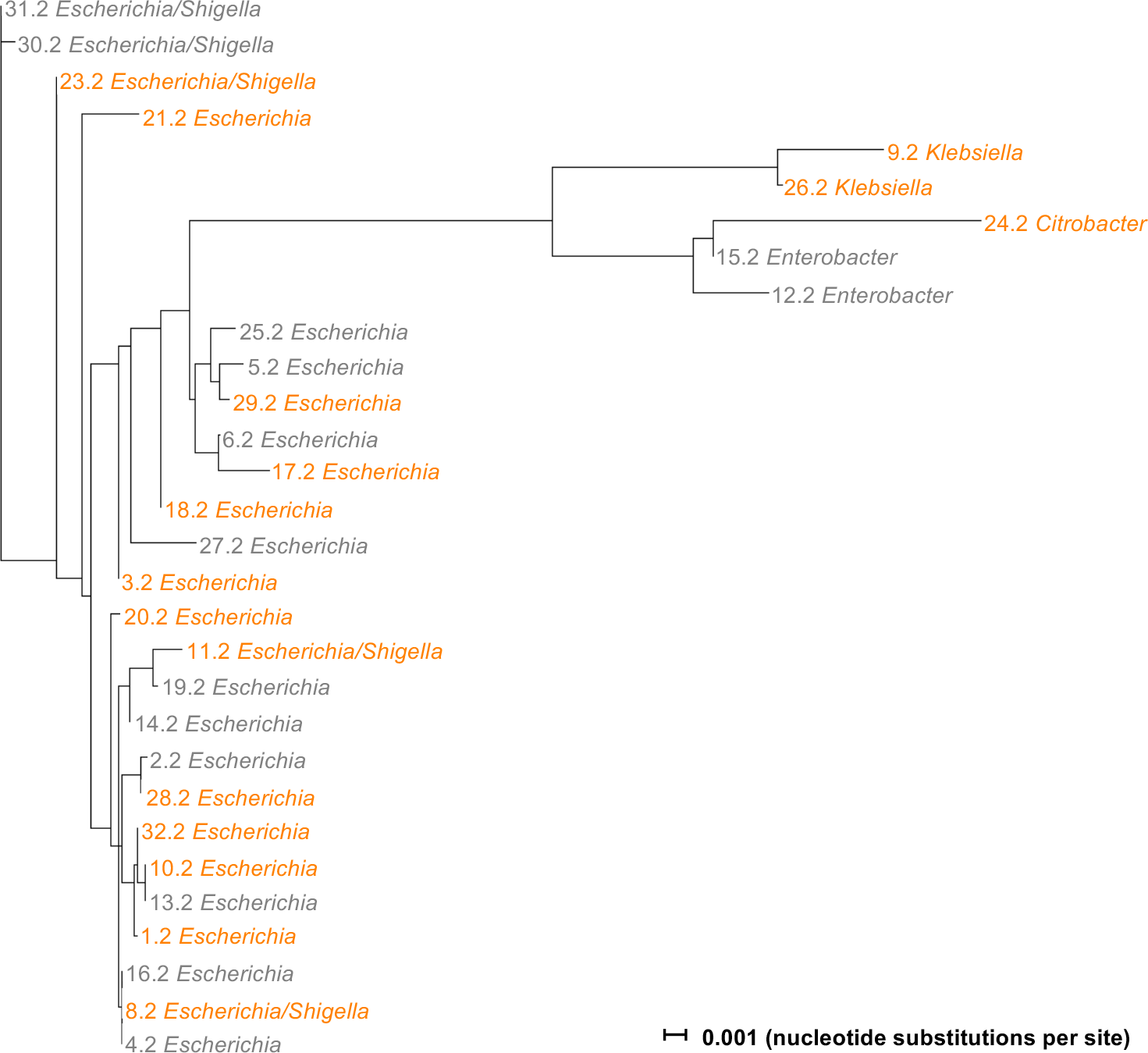
Phylogenetic tree of *Enterobacteriaceae* strains isolated one day after treatment. Tree showing the relatedness between bacterial strains (n = 30) isolated from participant samples one day after treatment with sucrose (grey) or lactulose (orange). 16S rRNA sequences were annotated by BLAST and aligned using MUSCLE and SplitsTree. *Escherichia/Shigella* thereby indicates a strain with a sequence compatible with both, *E. coli* and *Shigella*. Horizontal lines indicate nucleotide substitutions per site. Please note that *E. coli* and *Shigella* are extremely closely related. Thus, the detection of *Escherichia/Shigella*-like sequences does not imply that *Shigella* spp. were present in the stools of any of the participants.

### Antibiotic resistance

Annotated human bacterial isolates were tested for resistance against different antibiotics. Seven antibiotics (imipenem, cefepim, gentamicin, kanamycin, ciprofloxacin, and trimethoprim/ sulfamethoxazole) from 5 classes (carbapenems, cephalosporins, aminoglycosides, fluoroquinolone and folic acid synthesis inhibitors) were chosen according to known genetic determinants of resistance in *Enterobacteriaceae* to evaluate resistance status in environmental settings (35).

All bacterial strains tested were susceptible to imipenem, cefepime, and kanamycin (Table S2). Bacterial resistance against ampicillin was common (11 out of 30 strains, 37%) followed by sulfamethoxazole/trimethoprim (7 out of 30 strains, 23%), ciprofloxacin (2 out of 30 strains, 7%) and gentamicin (1 out of 30 strains, 3%). Five bacterial strains were resistant to two or more antibiotics (Table S2). No extended spectrum beta-lactamase producing strains were identified.

### Stability of microbiota upon lactulose exposure

We used 16S rRNA gene sequencing to analyze the taxonomic composition of the intestinal microbiota after lactulose challenge. Overall, we observed no significant difference in bacterial diversity between lactulose and sucrose treated individuals at day 1 and day 14 compared to pre-treatment conditions (Figure 7A). The microbiota composition was also highly similar between lactulose and sucrose treatment. An example for phylum level compositions is provided in Figure 7B. No significant shift in the taxonomic composition was detected at any taxonomic level (paired false discovery rate (FDR)-corrected Wilcox tests).

**Figure 7:**
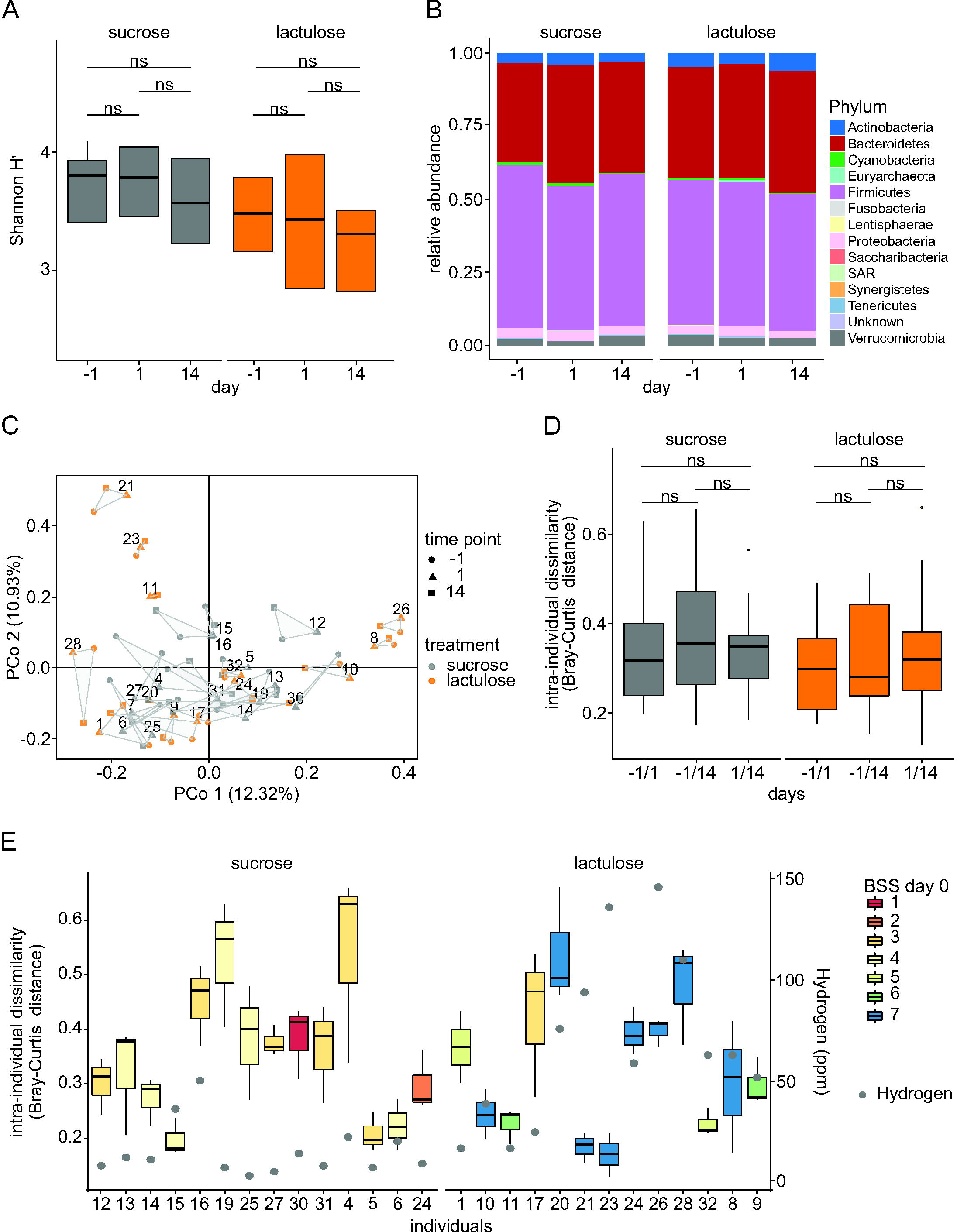
A single dose of lactulose (50 g) has no significant effect on fecal bacterial community structure a day or two weeks after treatment. (A) Shannon diversity index shows no significant change (paired Wilcoxon test, P>0.05) one day or two weeks after treatment (sucrose n=15, lactulose n=17) for each group. (B) Relative phylum abundance before (day −1) and after (day 1 and day 14) ingestion of sucrose (n=15) or lactulose (n=17). Colors indicate bacterial phyla, none on which were significantly different between time points. (C) Principal coordinate (PCo) analysis of microbiota composition with symbols and colors denoting sampling time point and treatment (sucrose, lactulose), respectively. The numbers denote the different individuals with samples from all three time point. (D) Intra-individual dissimilarity (Bray-Curtis) of microbial composition in samples collected at different sampling time points (day - 1 to 1, day −1 to 14 and day 1 to 14) and treatments with sucrose (n=14) and lactulose (n=13). Only individuals with samples from all three time points with a rarefaction cut-off of >19’000 were included in the intra-individual dissimilarity analysis (Bray-Curtis) of the microbiota composition. No significant shifts in bacterial composition were detected between different time points (paired Wilcoxon test, p>0.05). (E) Intra-individual dissimilarity (Bray-Curtis) of fecal microbial composition between the three sampling time points. The color key indicates the Bristol stool scale (BSS) of the same day (i.e., day 0) and after the lactulose/ sucrose challenge. The grey dots show maximal hydrogen levels in parts per million (ppm) following lactulose or sucrose ingestion.

We observed intra-individual shifts in the dissimilarity of microbial composition between the different sampling time points (days −1, 1, 14). Nevertheless, the shifts were similar in size for lactulose and sucrose treated individuals. Furthermore, we did not observe any clustering of samples by treatment (principal coordinate (PCo); analysis Figure 7C). Similarly, we detected no significant intra-individual dissimilarity changes before and after lactulose/ sucrose ingestion (Figure 7D). Additionally, there was no correlation between intra-individual dissimilarity (i.e. dissimilarity between time points) and BSS or H_2_ levels (Figure 7E).

Given the absence of a strong treatment-related effect on microbial composition, we sought to assess the sources of variation in our data set. To this end, we tested the relative contribution of individuality and treatment to the overall variation of microbial compositions (permutational MANOVA). For both treatments, we found individual variations but not treatment to explain the vast majority of the variation (effect of individuals: sucrose group - R^2^: 0.77, p<0.001; lactulose group - R^2^: 0.82, p<0.001; effect of treatment: sucrose group – R^2^: 0.01 n.s.; lactulose group – R^2^: 0.007, n.s.).

## Discussion

We performed a placebo controlled randomized controlled study to test effects of a single dose lactulose challenge on gastrointestinal microbiota composition and wellbeing in a homogenous group of young healthy individuals. The following key observations were made: i) Lactulose caused an increase in H_2_ levels in expiratory air and diarrhea in the vast majority of individuals in the lactulose group. ii) In the lactulose group, H_2_ levels correlated with severity of diarrhea (i.e. number of defecations). iii) Lactulose exposure did not increase the number of CFU of *E. coli* in stool samples collected 1 day after exposure. iv) The microbiota composition at that time point was also not significantly affected by single-dose lactulose challenge compared to the sucrose control, indicating that individual day-to-day variations of the microbiota are much stronger than the perturbation by a single 50 g lactulose ingestion.

### Effectiveness of a single dose lactulose as a laxative

Usage of lactulose for the treatment of acute or chronic constipation is well established (38). Recent randomized controlled trials used lactulose as a standard treatment for comparative testing against sodium polyethylene glycol (PEG) (39, 40), sodium picosulfate (39) and Chinese or Pakistan herbal medicine (41, 42). Effects of PEG were stronger than lactulose in a Cochrane meta-analysis (43) and PEG or liquid paraffin were more effective in children (44) even though these differences are small and possibly not clinically relevant. Cost-effectiveness of PEG and lactulose was virtually identical (45). Interestingly, lactitol might be similarly effective, but more palatable with lesser side effects than lactulose (46).

We are not aware of a recent placebo-controlled randomized controlled trial for one-time application of lactulose in young healthy individuals. Thus, our study provides valuable information on the effectiveness of lactulose in healthy individuals. We observed diarrhea (i.e. ≥3 bowel movements in 6 hours) in 14 out of 17 (82%) of healthy participants ingesting lactulose which was accompanied by at least moderate borborygmi, bloating, flatulence (14 out of 17, 82% for each symptom) and abdominal pain (8 out of 17, 47%).

### Methane levels in expiratory air and H_2_ non-producers

A fraction of patients and healthy individuals contain a methanogenic organism such as *Methanobrevibacter smithii* (47). Hydrogenotrophic methanogens use 4 molecules of H_2_ and 1 molecule of carbon dioxide (CO_2_) to produce 1 molecule of methane (CH_4_) (48). Due to this additional metabolization step, the CH_4_ peak will be delayed and due to the 4:1 ratio of H_2_ and CH_4_, the methane peak will be at a lower level compared to H_2_. Therefore, a cut-off of 10 ppm for CH_4_ is suggested (10). Furthermore, individuals with a methanogenic microbiota usually have increased levels of CH_4_ even at baseline, but fail to produce measurable H_2_ upon appropriate testing (10). In our analysis, methane measurements from only 14 individuals were available and only for one person, methane levels exceeded the threshold of 10 ppm. Our study included three H_2_ non-producers (18%) without any increase in H_2_ levels, well within the expected range of 2-43% in healthy individuals or patients (49). However, no methane measurements for these individuals are available.

### Correlation of symptoms with H_2_ levels

The healthy human intestine contains 30-200 ml (average 100 ml) gas, which mainly consists of H_2_, CO_2_ and CH_4_ (10, 50). Both, H_2_ and CH_4_ are exclusively produced by the bacterial microbiota (47).

Our study found a significant association of peak H_2_-levels with severity of diarrhea upon lactulose challenge in healthy individuals, arguing for a relationship between H_2_-production and symptoms after lactulose challenge. The association of results of the lactulose breath test with a diagnosis of irritable bowel syndrome (IBS) or IBS symptoms is controversial. A large analysis failed to find differences in IBS patients vs. healthy controls (51). However, the H_2_ response in the lactulose breath test was related to bloating in IBS patients (52–54). Similarly, a recent study using a large cohort of Chinese IBS patients found both, high H_2_ levels (area under the curve) in breath tests and visceral hypersensitivity to be risk factors for bloating and borborygmi and overall symptom burden (55).

Methane data were available only for a subset of participants and no correlation with symptoms could be established. Previous studies confirmed an association of methane levels in expiratory air with constipation (56, 57) and the severity of constipation correlated with methane levels (56).

### Stability of the microbiota upon single-dose lactulose challenge

In our study, no significant effects of a single dose of 50 g lactulose on microbiota composition could be detected after 1 day or after 14 days. Previous diet intervention studies showed that short-term macronutrient shifts reversibly alter gut microbiota composition within 24 hours even though broad patterns of microbiota composition similarities (enterotypes) remain stable (22–25). Our PCo analysis showed that lactulose challenge did not significantly alter the microbial composition since similar patterns of small-scale shifting were observed for lactulose and sucrose, alike (day 1 and day 14). Instead, our data argue for a stability of microbiota compositions with daily variations dominating over changes due to a single intervention with 50 g lactulose, even though diarrhea was observed in the majority of individuals. However, the data of this study provides no information about short-term microbiota compositional changes immediately after lactulose challenge, nor about site-specific effects within the small intestine or the right-sided colon.

### Effects of lactulose on the risks for intestinal infections

Prebiotics are food ingredients, which are neither digested nor degraded in the stomach or small intestine, but fermented by the gut microbiota, leading to a selective stimulation of the growth of certain intestinal bacteria and may thereby reap potential benefits for the host (58). Lactulose, along with inulin and other oligosaccharides are considered prebiotics and *Lactobacillus* and *Bifidobacterium* are considered significant target genera for the associated prebiotic effects (58, 59).

Since over 50 years, lactulose has been considered the “bifidus factor”, increasing fecal *Bifidobacterium* counts (60). A number of subsequent studies have confirmed increased intestinal content of *Bifidobacterium* (61–67) and *Lactobacillus* (61, 64) in human stool samples upon lactulose exposure. Those studies relied on conventional microbiological techniques and did not address the overall composition of the intestinal microbiota. Furthermore, in contrast to our study, lower dosages of lactulose (5-20 g) and longer exposure times (4–8 weeks) were used, potentially explaining differences to our results. In fact, effects on *Bifidobacterium* stool densities were weaker and not significantly differed from controls in an 8-day study (68). However, short-term effects of lactulose on the intestinal microbiota have not been tested previously. No “bifidus factor” effects were observed in our study and differences in the time of lactulose exposure might explain this discrepancy.

Fermentation of lactulose is not limited to *Lactobacillus* and *Bifidobacterium*. Glucosidases and bacterial transporters for lactulose are abundant in a large number of intestinal bacteria including *Cronobacter*, *Enterococcus*, *Escherichia*, *Klebsiella*, *Pseudomonas*, and *Streptoccoccus* spp., which were able to grow on lactulose as the sole carbon source (13). One recent study addressed effects of feeding mice with lactulose for two weeks: The authors noted a decrease in pH and an increase in short chain fatty acids in the colon. In the same mice, an increase in *Proteobacteriacae* and *Actinobacteria* including *Bifidobacterium* and a decrease in *Firmicutes* was observed. An increase in *Helicobacter* and *Akkermansia* spp. content was interpreted to be secondary to increased mucin production. Similar to our study, no increase in *E. coli* content was observed (69). These changes were accompanied by an increase in *Bifidobacteriae*, *Lactobacillus* spp. and *Enterobacteriacae*. In our study, we found a remarkable stability of the microbiota upon lactulose perturbation with 50 g lactulose resulting in diarrhea in >80% of participants. In fact, day to day variation in the control group were as big as variations in the experimental group and only approximately 3% of the variance can be explained by lactulose treatment (not shown).

Effects of lactulose on growth of intestinal pathogens are controversial. *In vitro* co-cultures of human intestinal *Lactobacillus* isolates with *E. coli*, *Salmonella enteritidis* or *Vibrio cholerae*, demonstrated reduced growth of the enteric bacterial pathogens (70). Case series also demonstrated therapeutic effects of lactulose treatment for human carriers of non-typhoid *Salmonellae* (71) and a temporary reduction of *Shigella* excretion in carriers (72). Application of lactulose was suggested to decrease the number of urinary tract infections (73). However, in a rat model of *Salmonella enteritides* infection, lactulose inhibited *S*. Typhimurium colonization, but stimulated bacterial translocation and intestinal inflammation (74).

In our study we were aiming to establish an experimental model for improved growth conditions for *Enterobacteriace. In vivo* data from our group with a murine colitis model suggested that the initial growth of *S*. Typhimurium within the first day depended on the presence of enzymes enabling hydrogen utilization (18). Better growth conditions for *S*. Typhimurium would suggest a higher susceptibility for bacterial infections upon lactulose exposure. This might be relevant for patients with liver cirrhosis which are highly susceptible to bacterial infections (75) and for whom lactulose remains the first-line drug for treatment and prevention of hepatic encephalopathy (9).

In our study, we did not find an increase in *E. coli* levels one day after lactulose application even though *E. coli* could potentially benefit from H_2_ produced upon lactulose ingestion. In agreement with our results, no clinical data support an increased risk of enterobacterial infections upon lactulose exposure. One reason for this could be that intrinsic *E. coli* are part of metabolic networks in biofilms in the large intestine and might still depend on metabolic contributions from other bacteria (according to the "Restaurant" hypothesis) (76), which limits any stimulatory effects of H_2_ and further studies for the metabolization of lactulose in the intestine are warranted.

In summary, exposure with a single lactulose dose leads to diarrhea in >80% of healthy individuals but not to an increase in *E. coli* CFU and no microbiota shifts. A single lactulose dose is therefore unlikely to significantly improve growth conditions of H_2_-consuming enteropathogens such as *S*. Typhimurium in the human gut.

## Author contributions

L. M., G. R., B. M. and W.-D. H. designed the clinical study. S. Y. W. and B. M. conducted the clinical study; S. Y. W., B. D., M. K., M.Z., S. S., and B. M. performed the experiments and analyzed the data. M. F., L. B., D.P., H.H., M. S. contributed to the analysis and interpretation of the data, S. Y. W., W.-D. H. and B. M. wrote the paper. All authors approved the final version of the manuscript.

## Supplementary material

**Supplemental Figure S1: Relationship of bacterial counts with hydrogen levels and clinical symptoms.** (A) Correlation analysis between maximum (max) hydrogen levels and bacterial loads within feces collected one day after treatment. (B) Correlation analysis between bacterial loads within feces collected one day after treatment and the number of defecations during 6 hours following lactulose or sucrose ingestion. (C) Correlation between bacterial loads within feces collected one day after treatment and Bristol stool scale during 6 hours following lactulose or sucrose ingestion. p > 0.05 = not significant; **p < 0.01 = significant; Spearman R correlation. Dashed lines = detection limit.

**Supplementary Table S1: Detailed quantification of symptoms by individual participants.**

**Supplementary Table S2: Antibiotic resistances of isolated *E. coli* strains.** Medium grey indicates resistance against the respective antibiotics, dark grey means medium resistance against the tested antibiotic (this depends on the antibiotic) and light grey means susceptible to the antibiotic.

## Abbreviations

BLAST: basic local alignment search tool
BSS: Bristol stool scale
CFU: colony forming units
CR: colonization resistance
LB: lysogeny broth
OD: optical density
PBS: phosphate buffered saline
PEG: Polyethylene glycol
SIBO: small intestinal bacterial overgrowth
*S*. Typhimurium: *Salmonella* Typhimurium

